# Characterisation of the RNA-Binding Properties of the MRSA β-lactam resistance enzyme PBP2a

**DOI:** 10.64898/2026.07.05.736576

**Authors:** Niki Christopoulou, Ngô Hồng Dương, Pedro Arede-Rei, Gabriel Torrens, Matthieu Blandenet, Felipe Cava, Sander Granneman

## Abstract

Analysis of RNA-binding proteome data from different bacterial species revealed many cell wall metabolic enzymes cross-linking to RNA *in vivo*, hinting that these proteins directly bind RNA. Surprisingly, penicillin-binding proteins (PBPs) were also abundantly identified as putative RNA-binding proteins. The cell surface localisation properties of many of these proteins therefore beg the question at what stage of their cellular life cycle these proteins interact with RNA and what the functional significance is. Here, we characterised the RNA-binding activity of PBP2a, the alternative transpeptidase that confers β-lactam resistance in MRSA. Using *in vivo* RNA-binding assays, we show that PBP2a interacts with hundreds of transcripts without apparent sequence specificity. Computational analyses identified a possible RNA-binding cleft in PBP2a proximal to its active site. Mutation of only two predicted positively charged residues located in this cleft substantially reduced cross-linking *in vivo*, implying that RNA recognition is largely dictated by RNA backbone interactions. While PBP2a does not regulate RNA steady-state levels, RNA-binding appears important for proper protein function: an RNA-binding deficient mutant exhibits reduced oxacillin resistance. These findings establish PBP2a as an RNA-binding protein *in vivo* and provide a framework to investigate how this non-canonical interaction may relate to cell wall biogenesis and β-lactam resistance.

## Introduction

Methicillin-resistant *Staphylococcus aureus* (MRSA) is an important human and animal pathogen causing healthcare problems worldwide. It is renowned for its capacity to rapidly adapt to the host environment and alter its cellular functions to evade the host immune system and establish successful infections. Swift alterations in gene expression are key to these rapid adaptive responses, with RNA-binding proteins (RBPs) playing an important role in post-transcriptional regulation of gene expression. In recent years, more studies have focused on the identification of RBPs in pathogens, shedding light on the contribution of RBPs to bacterial adaptation and infection. With the rapid development of high-throughput techniques that allow global capture of RNA-binding proteomes (RBPomes), recent studies have unearthed a plethora of novel RBPs in bacteria (1–5). These studies also predicted that many metabolic enzymes could bind RNA, suggesting a link between RNA regulation and metabolism. However, many of these predictions remain to be validated.

Previous work in our lab uncovered the RBPome of two clinically relevant MRSA strains, revealing hundreds of new putative RBPs (2). Much to our surprise, enzymes involved in the cell wall biosynthesis pathway were among the isolated RNA-binding proteins, several of which were also detected in previous RNA-binding proteome analyses on Gram-negative bacteria (1, 4). Interestingly, among these cell wall synthesis enzymes were penicillin-binding proteins (PBPs; Figure 1A). PBPs are transmembrane enzymes facing the periplasmic side of the cell wall. They are responsible for the final steps of peptidoglycan synthesis, the main component of the bacterial cell wall. Peptidoglycan consists of alternating N-acetylglucosamine (NAG) and N-acetylmuramic acid (NAM) sugar units cross-linked by short peptide chains. It is synthesised through glycan chain polymerisation by transglycosylases and subsequent cross-linking of peptide side chains by transpeptidases (TPases). *S. aureus* has four native PBPs (PBP1-4), which have transpeptidase and/or transglycosylase activity. However, MRSA strains have acquired an extra PBP, named PBP2a, which has low affinity for these antibiotics, leading to high levels of β-lactam resistance and therefore rendering MRSA infections treatment extremely challenging (6, 7). Notably, ceftaroline, a fifth-generation cephalosporin, retains activity against MRSA by binding covalently to PBP2a (reviewed in (8)). When β-lactams fail, vancomycin, a glycopeptide antibiotic that inhibits peptidoglycan synthesis through a different mechanism by binding to the D-Ala-D-Ala terminus of peptide precursors, remains a critical treatment option for severe MRSA infections.

**Figure 1:**
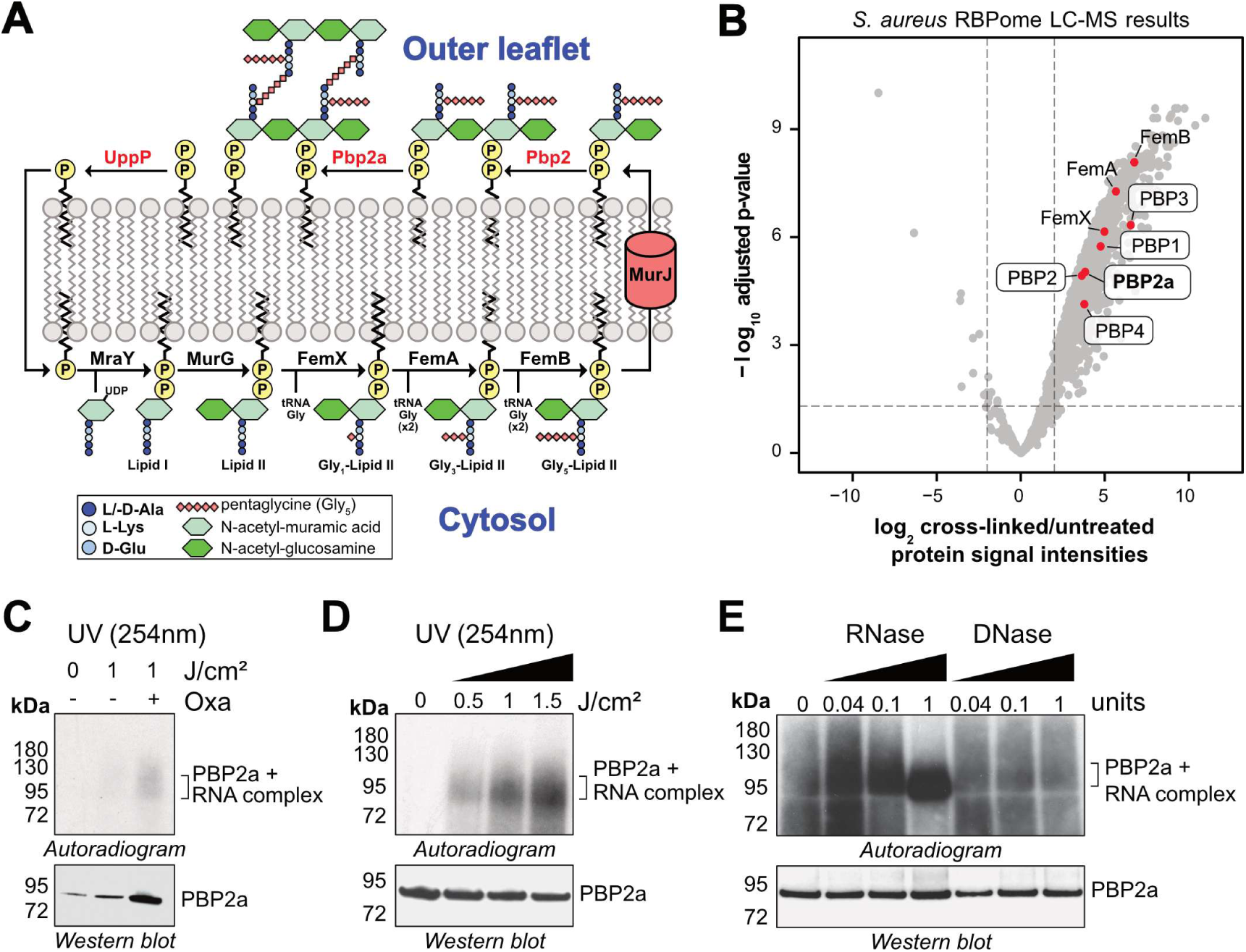
PBP2a binds RNA *in vivo*. **A)** Peptidoglycan synthesis in *S. aureus* (Adapted from (15, 16)) commences in the cytoplasm, where the MurA, MurB, MurC, MurD, MurE, MurF enzymes catalyse a series of reactions converting UDP-GlcNAc to UDP-MurNAc attached to the stem peptide. MraY binds UDP-MurNAc-pentapeptide to the membrane associated C55-PP, generating lipid-I, which is then converted to lipid-II, with the addition of a UDP-GlcNAc, mediated by MurG. The Fem proteins attach five glycine residues to the lysine of lipid-II and then the occurring Gly5-lipid-II is exported by MurJ to the cell surface. It is then polymerised by transglycosylases, like PBP2, and cross-linked by transpeptidases, like PBP2a, PBP2 and other native PBPs. While β-lactams inhibit the TPase activity of native *S. aureus* PBPs, PBP2a has low affinity for most β-lactam antibiotics, enabling peptidoglycan cross-linking. **B)** Peptidoglycan synthesis enzymes enriched in the RBPome of MRSA. The protein signal intensity values obtained with LC-MS were compared between cross-linked and untreated (negative control) samples. **C-E**) PBP2a cross-links to RNA *in vivo*. After UV cross-linking (254nm) of *mecA*-HTF cells, PBP2a-RNA complexes were purified under stringent and denaturing conditions (2). Following RNase treatment and RNA radiolabelling, samples were resolved on NuPAGE gels and PBP2a-RNA complexes were detected by autoradiography. The protein (82 kDa) was detected by Western blotting using an anti-TAP antibody. **C:** Comparing the RNA cross-linking signals in the presence and absence of oxacillin (0.2 µg/ml). **D:** RNA cross-linking increased proportionally to the UV dose (J/cm^2^) used. **E:** Treatment of purified PBP2a with increasing amounts of RNase A/T1 or RQ1 DNase. Cells were grown in the presence of 0.2 mg/L oxacillin and irradiated with 1.5 J/cm2 UV energy.

An obvious question is the biological significance of PBP-RNA interactions. Given their cell surface localisation, PBPs are not expected to encounter RNA as readily as cytosolic proteins, making such interactions inherently surprising. One possibility is that PBPs interact with extracellular RNA (eRNA), which is abundant in biofilms and could provide a functional interface for these enzymes (9, 10). Alternatively, RNA binding may occur earlier during protein biogenesis or transiently within the cell before membrane insertion. To address these possibilities and define the mechanistic basis of this interaction, we characterised the RNA-binding properties of *S. aureus* PBP2a both *in vivo* and *in vitro*.

## Results

### PBP2a binds RNA *in vivo* in *S. aureus*

Our previous RNA-binding proteome (RBPome) analyses of *S. aureus* strains utilised UV cross-linking combined with silica resin to purify total RNA as well as proteins that were covalently cross-linked to RNA by UV irradiation. This revealed hundreds of new putative RNA-binding proteins (RBPs), including many involved in cell wall metabolism (2). The Fem proteins, enzymes that use tRNA-Glycine to build the peptidoglycan pentapeptide bridge, were highly enriched in our RBPome data, as might be expected (Figure 1A–B). Much to our surprise, however, the PBP2a enzyme that confers resistance to β-lactam antibiotics, together with all four native *S. aureus* PBPs (PBP1–4), were similarly highly enriched (Figure 1B). These data therefore imply that PBPs have affinity for RNA. Given the critical role of PBP2a in resisting antibiotics, we validated the RNA-binding protein capacity of PBP2a *in vivo* and *in vitro*. We generated an *S. aureus* strain that expresses PBP2a fused to a His_6_-TEV-3xFLAG (HTF) tag at the C-terminus to perform *in vivo* RNA-binding experiments using the CLIP-related CRAC approach (11, 12). CRAC combines UV cross-linking followed by affinity purification of the tagged protein and RNA sequencing to identify direct protein-RNA interactions (see Methods). For our validation experiments, we chose to use the hospital-associated N315 strain (WT), as it is easier to manipulate genetically and we were able to successfully generate both a *mecA* knock-out (gene encoding PBP2a) and a PBP2a-HTF strain.

Previous work, however, had demonstrated that fusing fluorescent reporters to either N- or C-terminus of PBP2a lost their resistance to β-lactam antibiotics, suggesting that the fusion proteins have lost their enzymatic activity (13). Therefore, we tested whether the HTF tag would interfere with PBP2a function by assessing whether the C-terminal HTF tag impacted N315 resistance to the β-lactam oxacillin, as a proxy for PBP2a activity. Growth curve analyses in the presence of a concentration range of oxacillin showed that the growth of the strain expressing PBP2a-HTF was comparable to the WT strain (Supplementary Figure 1A), suggesting that the tag does not substantially impact PBP2a enzymatic activity.

Before starting the CRAC experiments, we also determined the optimal culture density where PBP2a was expressed at its highest level using Western blotting with anti-FLAG antibodies. These analyses were performed in TSB as well as TSB supplemented with oxacillin (0.2 µg/mL), to fully release the repression of the *mecA* gene encoding PBP2a. This indicated that PBP2a expression was generally highest during late exponential phase in both conditions (Supplementary Figure 1B-C; between OD_595_ of 3.0 and 5.0). Given that higher culture densities are more practical when performing CRAC experiments, an OD_595_ of 3.0 was used for mapping the RNAs that interact with PBP2a.

During the optimisation of the PBP2a-HTF CRAC experiments, we first determined the optimal UV 254nm cross-linking doses and conditions needed to detect significant PBP2a-RNA cross-linking. For this purpose, cells were grown in TSB or TSB supplemented with oxacillin to the desired OD_595_, followed by UV irradiation. Non-irradiated samples were used as negative controls. Following tandem affinity purification of PBP2a under highly denaturing conditions and partial RNA digestion, cross-linked RNAs were radiolabelled, and complexes were resolved by electrophoresis in NuPAGE gels. Radiolabelled RNAs were subsequently detected by autoradiography and PBP2a by Western blotting. The presence of a radioactive signal above the molecular weight of PBP2a provided evidence for RNA cross-linking. This was most apparent when N315 cells were treated with oxacillin, which boosted PBP2a expression (Figure 1C). The intensity of the radioactive signal was also proportional to the UV intensity dose used for the cross-linking (Figure 1D).

High doses of UV energy can also lead to cross-linking of proteins to DNA (14). For this reason, we performed additional CRAC experiments where we treated the *in vivo* cross-linked PBP2a-RNA complexes with a concentration gradient of RNase or DNase. If the radioactive signal originated from cross-linking to RNA, incubating the purified complexes with increasing RNase concentrations should lead to higher RNA degradation, thereby condensing the radioactive signal near the protein’s molecular weight. Consistent with this, the radioactive signal became more compact in the region of the gel where PBP2a migrates when higher RNase amounts were used (Figure 1E), indicating that the treatment shortened the cross-linked nucleic acids. In contrast, the signal remained unchanged and highly diffuse when samples were treated with increasing concentrations of DNase. This indicates that PBP2a preferentially UV cross-links to RNA in *S. aureus* cells.

After confirming the RNA-binding activity of PBP2a *in vivo*, we next isolated the cross-linked RNAs and converted them into next-generation sequencing libraries to investigate which RNAs interact with PBP2a (see Methods). To account for background RNA signal, we used UV cross-linked cells of the WT *S. aureus* strain as a negative control, which expresses untagged PBP2a. Three independent biological CRAC experiments were performed, each consisting of three technical replicates. All CRAC experiments produced very low background, as judged by the remarkably low number of reads detected in negative control samples (Supplementary Figure 2A–B). Comparison between biological replicates also showed that the read coverages for genes detected in the data were highly correlated (Supplementary Figure 2C; Pearson correlation coefficient [r] ≥ 0.8), indicating that the individual CRAC experiments generated reproducible results. Normalised transcript per million (TPM) read counts for each separate technical replicate and their distribution were comparable (Supplementary Figure 2D). Because the technical replicates for each sample had comparable read coverages, they were merged to generate three individual biological replicate samples.

Meta-analysis of TPM-normalised CRAC data revealed that tRNA and rRNA were the most abundantly detected RNA classes, with mRNAs being underrepresented (Figure 2A). We next asked whether PBP2a preferentially binds specific transcript regions (for example, 5′ or 3′ ends). Visual inspection of the data (bedgraph files, see Data Availability) showed that PBP2a bound many transcripts throughout their whole sequences. Focussing on tRNA and mRNAs, as representative examples, we analysed the global distribution of CRAC reads using the pyBinCollector script from the pyCRAC package (17). Briefly, this tool divided all mRNA transcripts that are at least 100 nucleotides long into 100 bins and tRNA into 50 bins to normalise for transcript length. It then counts how frequently reads were mapped to each bin and then sums these values for all genes (Figure 2B). The coverage plots produced for tRNAs (most highly recovered class), and mRNAs revealed a mostly random distribution of reads across transcript length (Figure 2B), implying that PBP2a does not prefer binding specific nucleotide sequences or gene regions. Some enrichment of PBP2a near the 5’ region of tRNAs was apparent (Figure 2B). However, visual inspection of the read coverage in bedgraph files on individual tRNAs from PBP2a-HTF and the WT negative control samples revealed similar binding profiles. As the data did not show a strong indication of specific binding of PBP2a to tRNA regions, we cannot rule out the possibility that some of these signals are background noise.

**Figure 2:**
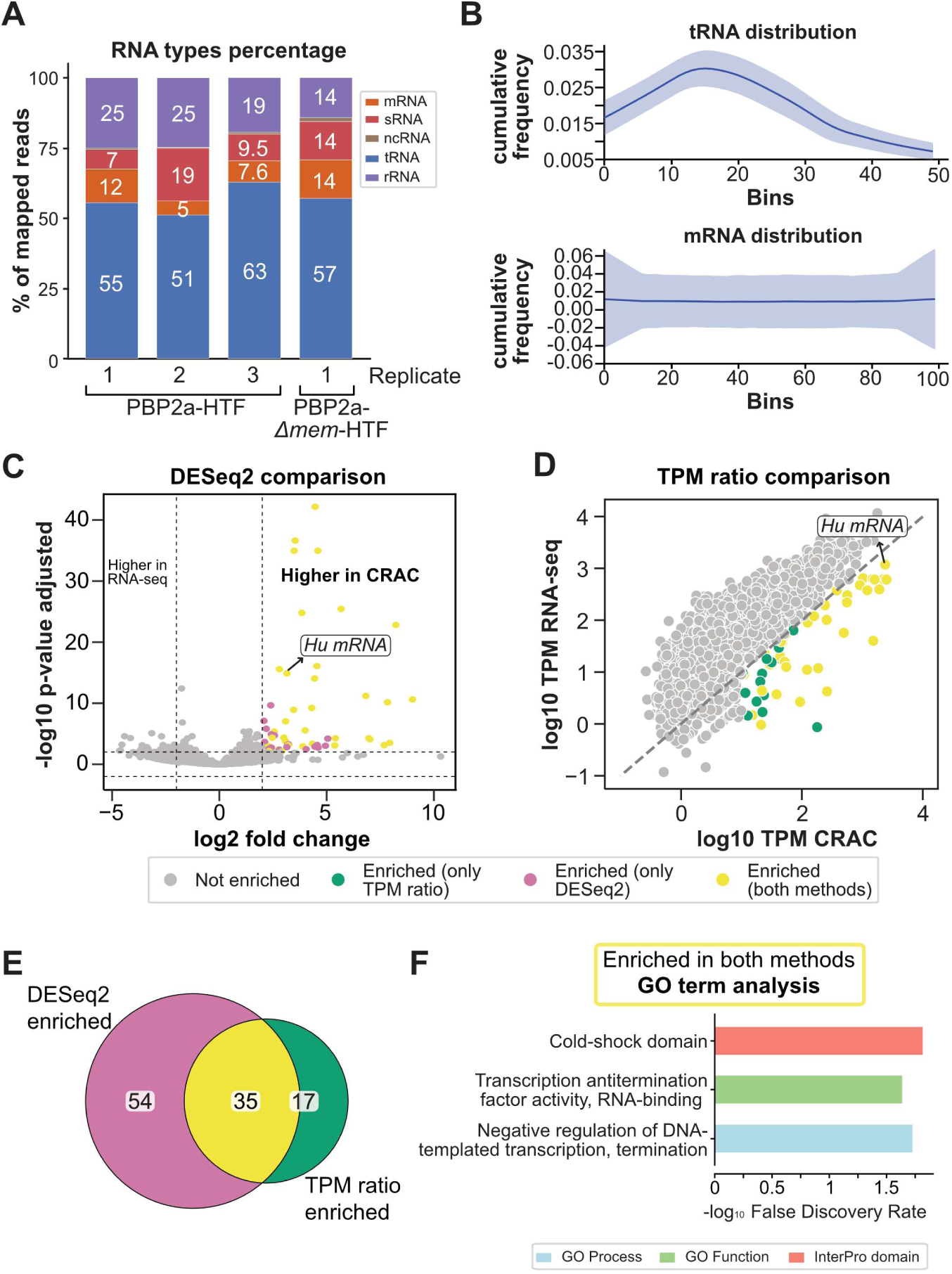
PBP2a binds hundreds of transcripts, with diverse sequences and functions. **A)** Representation of different RNA types in each biological replicate. TPM-normalised reads mapped in each RNA category were divided by the total number of TPM-normalised reads in each replicate. **B)** Read coverage over tRNAs and mRNAs normalised into 50 and 100 bins, respectively. **C)** Volcano plot presenting DESeq2 results. Coloured genes in the top right panel show significant enrichment in CRAC compared to RNA-seq. Yellow coloured dots represent transcripts that are also enriched in the TPM comparison method. **D)** Comparison of log10 TPM CRAC values, on the x-axis, with RNA-seq, on the y-axis. Coloured points represent transcripts enriched in CRAC relative to RNA-seq, with yellow indicating significance in both this method and DESeq2. **E)** Intersection of transcripts enriched in CRAC compared to RNA-seq from the two different methods presented in C and D. **F)** Gene ontology (GO) enrichment analysis of the 35 enriched transcripts was performed on STRING database. Shown is the False Discovery Rate (FDR) of the highest enriched GO terms.

We next compared our PBP2a CRAC data to RNA-seq reads generated from the same cross-linked cell cultures. We followed two different, but complementary, approaches to identify transcripts overrepresented in the CRAC data (**Error! Reference source not found.**C-D, see Methods). The first method was the DESeq2 approach, a tool designed to identify differentially expressed genes in RNA-seq data (18). Although this tool is often used to make such comparisons and has enabled us to detect genes seemingly enriched in the CRAC data (Figure 2C), the tool does assume that most genes are not differentially expressed. This could be problematic when comparing CLIP/CRAC directly to RNA-seq data, two vastly different methods that are very likely to show big differences in gene read counts. Therefore, we also performed a simpler analysis where we used scatter plots to compare CLIP and RNA-seq data, and identified transcripts enriched in the CLIP data by applying a fold-change threshold (Figure 2D). We then intersected the results and identified a set of 35 mRNAs enriched in the CRAC (**Error! Reference source not found.**E). Functional enrichment analysis of the proteins encoded by these mRNAs revealed enrichment of cold-shock proteins (**Error! Reference source not found.**F).

RBPs generally act upon the RNAs they bind by altering their stability, structure and/or translation. To determine whether PBP2a could have a role in regulating RNA steady-state levels, we examined the effects of PBP2a deletion on the transcriptome. For this, we compared the steady-state RNA levels from WT cells and N315 lacking PBP2a (*ΔmecA*). Analysis of the sequencing data (three independent replicate samples for each strain) did not reveal any transcript differentially expressed in the deletion mutant (except for the deleted *mecA* transcript that encodes PBP2a; Supplementary Figure 3A). PBP2a was expressed under the growth conditions used (Supplementary Figure 3B). Prior to receiving the RNA-seq data, we had already performed RT-qPCR analyses on selected transcripts (Supplementary Figure 3C) that showed robust PBP2a cross-linking in the CRAC data. This revealed that the absence of PBP2a had no significant impact on their RNA steady-state levels (Supplementary Figure 3C), which was confirmed by the RNA-seq data (Supplementary Figure 3D).

Collectively, these data imply that PBP2a binds RNA *in vivo* but does not directly impact the stability of RNAs it binds.

### PBP2a binds nucleic acids *in vitro*

To complement our analyses, we subsequently investigated the RNA-binding capacity of recombinant PBP2a (Supplementary Figure 4A) *in vitro*, using Electrophoretic Mobility Shift Assays (EMSAs). Based on the CRAC data, we designed RNA oligonucleotides, focusing on transcript regions that showed strong cross-linking in the CRAC data relative to the RNA-seq data (Figure 3). This included a non-coding RNA (Teg104; Figure 3A and 3C) and a fragment of an mRNA originating from the *SA_RS07385* gene that encodes for the DNA-binding protein Hu (Figure 3B and D). The EMSA results show that PBP2a was able to bind both RNA fragments (Figure 3E-F), with an estimated K_D_ of 1.5-2 µM (Figure 3G-H), suggesting moderate binding affinities. No binding of these RNAs was detected by our BSA negative control protein (Figure 3I-J). PBP2a complex formation with the DNA sequences of these RNAs could also be readily detected (Supplementary Figure 4B), although K_D_s could not be reliably estimated, given that the complexes resolved poorly in the gel. Closer inspection of the EMSA results (Figure 3E-F) showed that the band representing the PBP2a-RNA complex progressively migrated higher up into the gel as the concentration of PBP2a increased. We interpret this as oligomerisation of PBP2a on the RNA, possibly through cooperative binding. Therefore, the estimated K_D_ value should be interpreted with caution.

**Figure 3:**
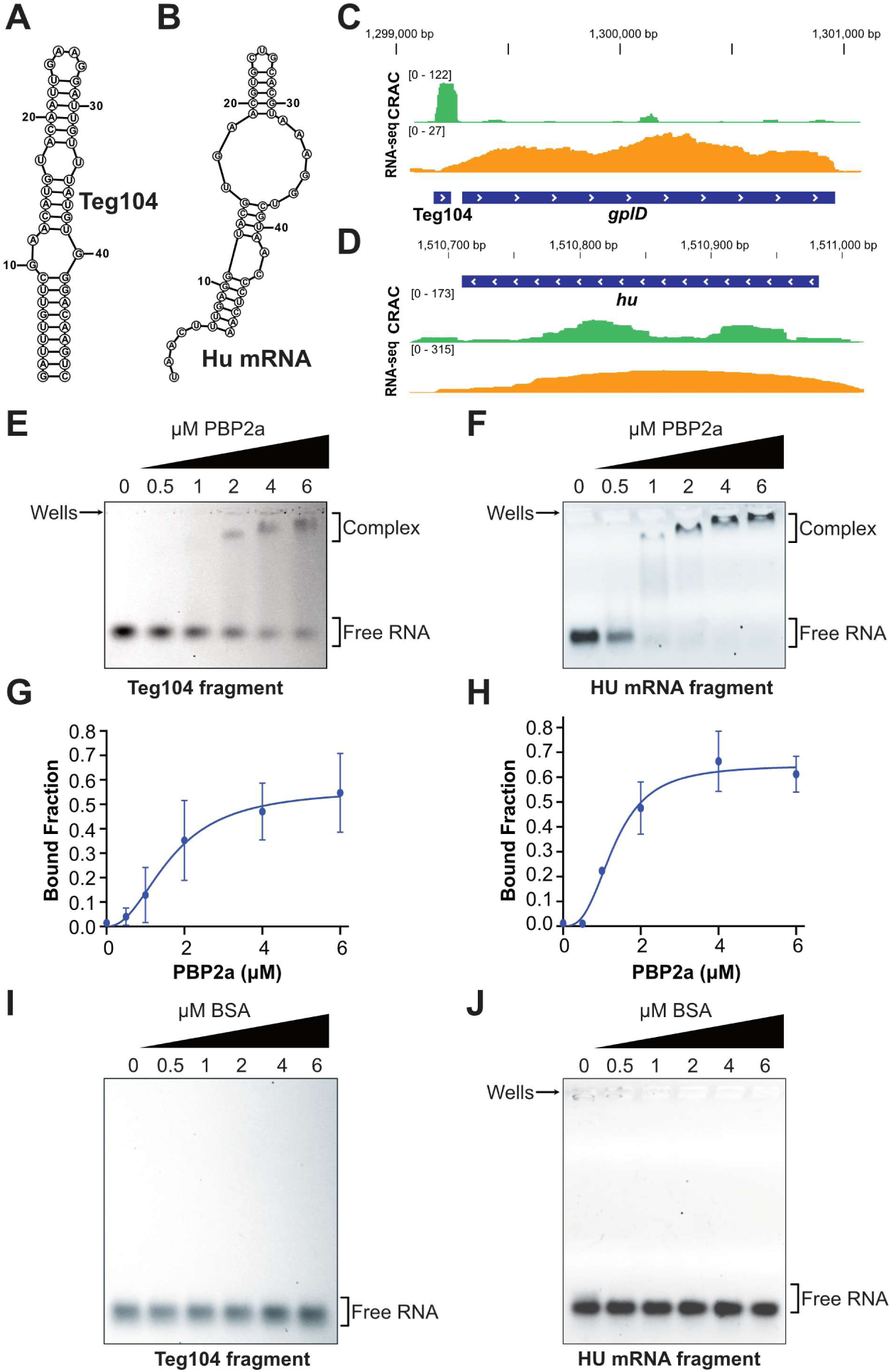
PBP2a binds RNA *in vitro*. **A-B**) Secondary structure of IRD-800-labeled Teg104 and 6FAM-labelled Hu RNA oligonucleotides selected for EMSA. The secondary structure was predicted using the RNAstructure tool (19). **C-D**) Genome browser snapshot of CRAC and RNA-seq sequencing reads mapped on the regions of the RNA oligonucleotides (visualised with Integrated Genomics Viewer (20)). **D-E**) EMSA performed with increasing amounts of PBP2a in the presence of 0.05 μM of 6FAM- or IRD800-labelled RNA fragments. **F-G**) Quantification of the EMSA results using ImageQuantTL v8.2.0 using data from two independent replicates. The bound fraction was calculated by dividing the signal intensity of the PBP2a-bound RNA with the total intensity of bound and free RNA. Binding curves were generated in GraphPad Prism v.8.0.1 using the Hill slope Least squares fit method. Plotted are the best-fit models, with mean values (round points) and standard deviations (error bars) highlighted. **H-I**) EMSA performed with increasing amounts of the negative control protein (BSA) in the presence of 0.05 μM of 6FAM- or IRD800-labelled RNA fragments.

### Forcing PBP2a to the cytosol increases RNA-cross-linking

PBP2a is localised at the cytoplasmic membrane and extends into the extracellular space where it participates in cell wall synthesis, anchored in the membrane (21). Therefore, we were puzzled that it was detected in our RNA-binding proteome analyses. While extracellular RNA (eRNA) has been detected in *S. aureus* biofilms, it seemed less likely for PBP2a to bind RNA in the cell wall environment, as the RNA concentration is presumably much lower compared to the cytosol. Moreover, the current assumption is that the unfolded nascent PBP2a polypeptide is translocated by the general secretion machinery where it is folded in the extracellular environment (22). However, it is unclear whether PBP2a is co-translationally translocated or only once it is fully synthesised. Thus, there may be a possibility that RNA and PBP2a interact in the cytosol, although in this scenario, RNA would likely encounter PBP2a in a largely unfolded state. To test whether PBP2a can interact with RNA in the cytosol, we constructed a strain that expresses a version of PBP2a lacking the transmembrane anchor (*Δmem*; amino acids 7-24). Constructs expressing HTF-tagged PBP2a alleles from its endogenous promoter were then introduced into the *ΔmecA* strain. Before determining the RNA-binding activity of the *Δmem* mutant, we first assessed the growth phenotype of the mutant strain. Deleting the PBP2a transmembrane anchor did not cause a notable growth defect under normal growth conditions in TSB, nor did it noticeably affect the expression levels of the protein (Figures 4A-B). However, the *Δmem* mutant exhibited oxacillin resistance levels similar to the deletion mutant (Figure 4A: -/- and empty plasmid plots). These results are consistent with PBP2a failing to localise to its site of action.

**Figure 4:**
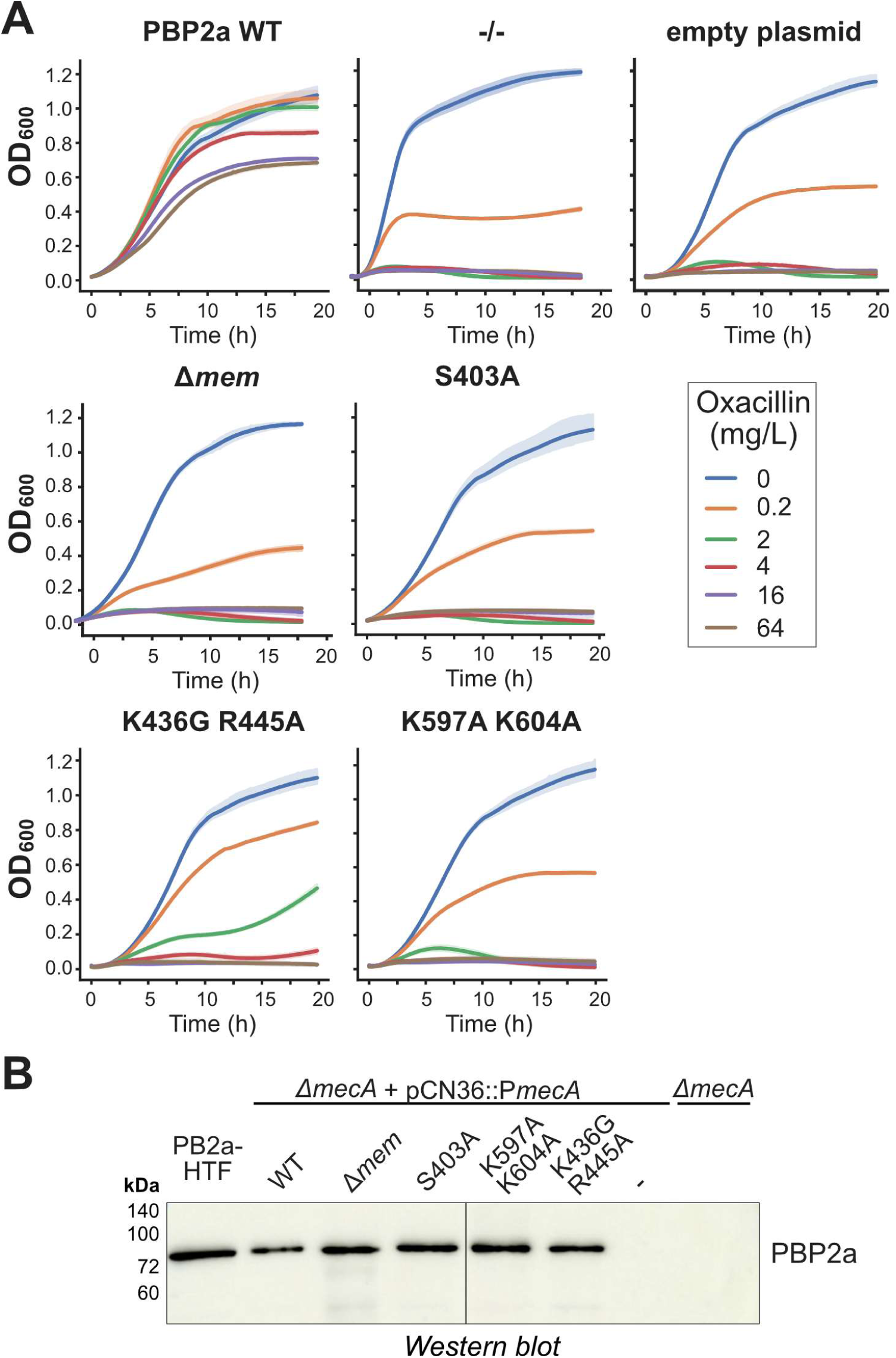
PBP2a mutations that reduce RNA-binding *in vivo* reduce oxacillin resistance. **A)** Growth curve analysis of the N315 *mecA* deletion strains complemented with wild-type PBP2a proteins or the indicated PBP2a mutants in the presence or absence of oxacillin. -/- indicates no plasmid and empty plasmid is deletion strain plus empty pCN36 plasmid. Each colour shows the results from a different concentration of oxacillin (µg/mL). **B)** Western blot analysis of the N315 strain expressing the genomically encoded PBP2a-HTF strain to the *ΔmecA* deletion strain complemented with the indicated plasmids expressing WT and mutant HTF-tagged PBP2a proteins. Western blot analysis was performed using anti-FLAG antibodies.

CRAC experiments revealed that the *Δmem* mutant UV cross-linked ∼7 times more efficiently to RNA relative to the wild-type counterpart (Figures 5A-B). Analysis of the cross-linked sequences revealed that the *Δmem* mutant CRAC data were highly correlated with the CRAC data generated with PBP2a-HTF (Supplementary Figure 3B-D). Again, tRNAs were the most enriched class of transcripts (Figure 2A). Collectively, these results show that PBP2a, even when it is likely not in its final folded state (22), can bind RNA in the cytosol.

**Figure 5:**
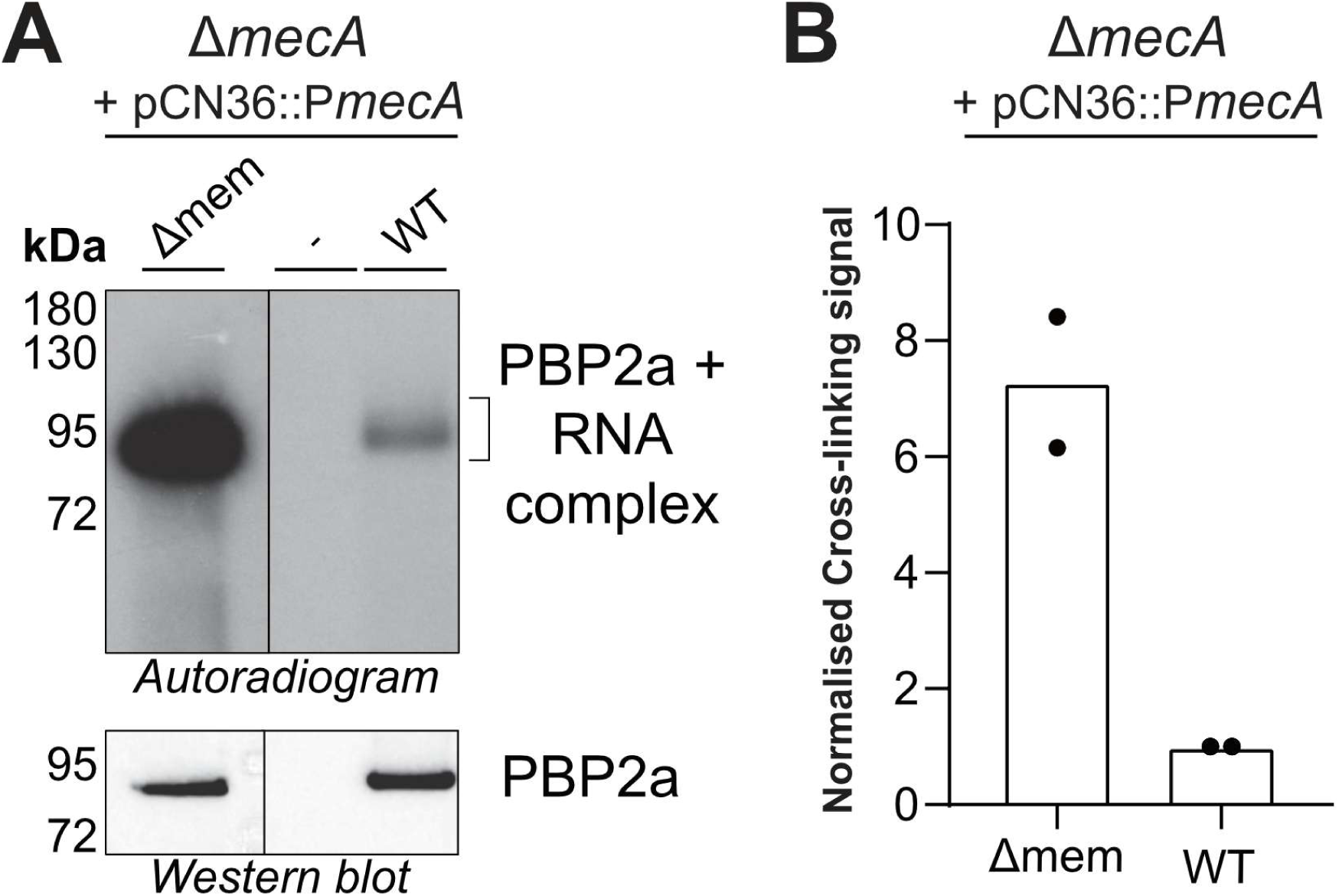
Deletion of PBP2a transmembrane anchor increases PBP2a cross-linking to RNA. **A)** CRAC analysis of the wild-type and *Δmem* PBP2a mutant. The top panel shows the autoradiography signal, indicating cross-linking of PBP2a to RNA. The bottom panel shows the Western blot analysis of the proteins purified during CRAC. **B)** Quantification of the relative cross-linking changes between the mutant and wild-type PBP2a, the cross-linking signals were normalised to protein levels, determined by western blotting, and then compared to the signal of the wild-type PBP2a, which served as a positive control. ImageQuantTL v8.2.0 was used for quantifying autoradiography and Western blot signals.

### PBP2a binds RNA near the transpeptidase domain

Having established that PBP2a binds RNA *in vitro* and *in vivo*, we next determined which amino acids in PBP2a were responsible for this activity. For this purpose, we analysed the crystal structure of PBP2a (1VVQ) using our pyRBDome pipeline (23). Briefly, this Python pipeline submits the sequence and/or structure to five independent tools that predict whether amino acids within the protein can bind RNA and/or have pockets that are suitable for binding other ligands. The analyses include results from PST-PRNA (24), that predicts RNA-binding residues based on structure (topology) and sequence conservation, BindUP (25), a tool that identifies positively charged patches on the surface of protein structures (but does not consider sequence conservation), RNABindRPlus, a method that only uses sequence information to predict RNA-binding residues (26) and finally DisoRDPBind, a method that predicts RNA-binding residues in disordered protein regions (27). We then used the outputs of these predictors to train an XGBoost ensemble model (referred to hereafter as pyRBDome model) that uses these predictions to come up with a final RNA-binding probability for each residue. The reason for choosing this specific pipeline was because it was designed to help with predicting RNA-binding regions in non-canonical RNA-binding proteins, such as PBP2a (23). The results of individual prediction algorithms and the final predictions of the pyRBDome model are shown in Figure 6A-D. Our pyRBDome analysis identified a cleft within the PBP2a structure with elevated RNA-binding probability (Figure 6D). To substantiate these predictions, we performed molecular docking studies with short RNA oligonucleotide sequences using HDOCK (28). This confirmed the presence of favourable RNA-binding interactions within this cleft (Figure 6E). Finally, AlphaFold3 modelling with PBP2a and an RNA sequence that strongly cross-linked to PBP2a in the CRAC data (Teg104 sRNA fragment; Figure 3A) also implied that this region accommodates RNA binding in a manner consistent with the pyRBDome predictions (Figure 6F). Note that AlphaFold3 could not confidently predict the structure of the RNA in this complex (Supplementary Figure 5A-B).

**Figure 6:**
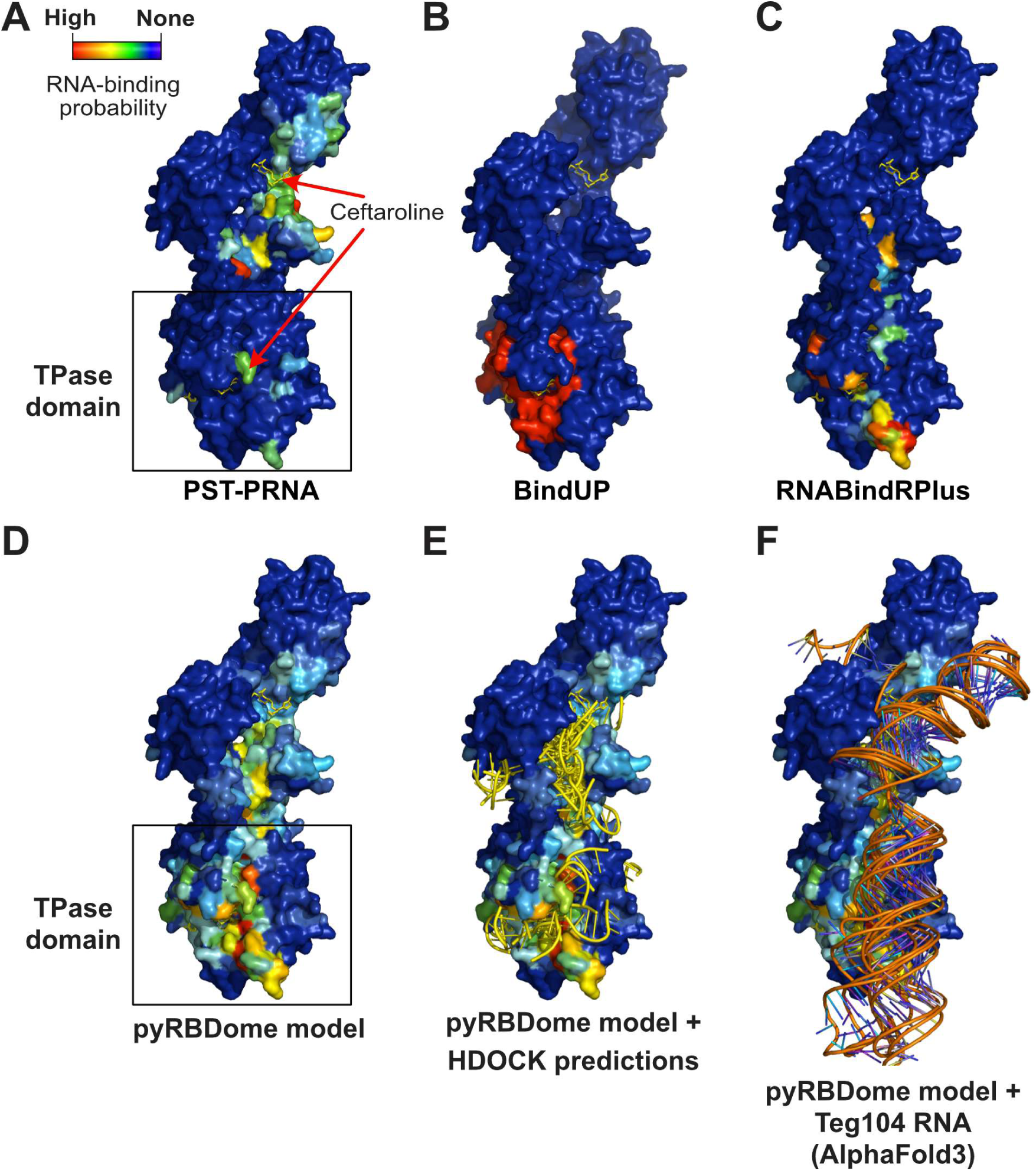
*In silico* analysis predicts the presence of an RNA-binding cleft in PBP2a. **A-C**). Results of PST-PRNA, BindUP and RNABindRPlus predictions using the PBP2a structure (rcsb ID 3ZFZ). PBP2a consists of the transpeptidase domain (TP) and the allosteric domain (top part). Two molecules of β-lactam antibiotic ceftaroline are bound on the allosteric and the active site of the transpeptidase domain in this structure (yellow sticks). The colours indicate RNA-binding probability. Cooler shades (blue) indicate lower, while warmer shades (red) show higher RNA-binding probabilities. **D**) The pyRBDome model represents a combination of all algorithms. **E-F**) pyRBDome prediction results superimposed on HDOCK prediction results (28) with short pyrimidine oligonucleotides (**E**) and with AlphaFold3 prediction results (29) showing PBP2a in complex with a Teg104 sRNA fragment.

### Mutating two positively charged residues in the predicted RNA-binding region significantly reduces PBP2a RNA-binding

To further dissect the RNA-binding activity of PBP2a, we used the pyRBDome predictions to design mutations that could disrupt RNA binding. We focused on mutating positively charged residues (e.g., arginine and lysine) located on the surface of PBP2a. We targeted residues with medium to high RNA-binding probabilities (shown in green to yellow; R445, K436, K597, and K604; Figure 7A) as well as residues with very low RNA-binding probabilities (K662, K663, K176, and K181), which should not impact RNA binding if mutated. In each PBP2a mutant, two proximal amino acids were simultaneously mutated to increase the likelihood of disrupting RNA binding. Finally, we also included the S403A mutant, which renders the transpeptidase catalytically inactive. These mutants were cloned into pCN36::P*mecA* with an HTF tag and introduced into the *ΔmecA* strain.

**Figure 7:**
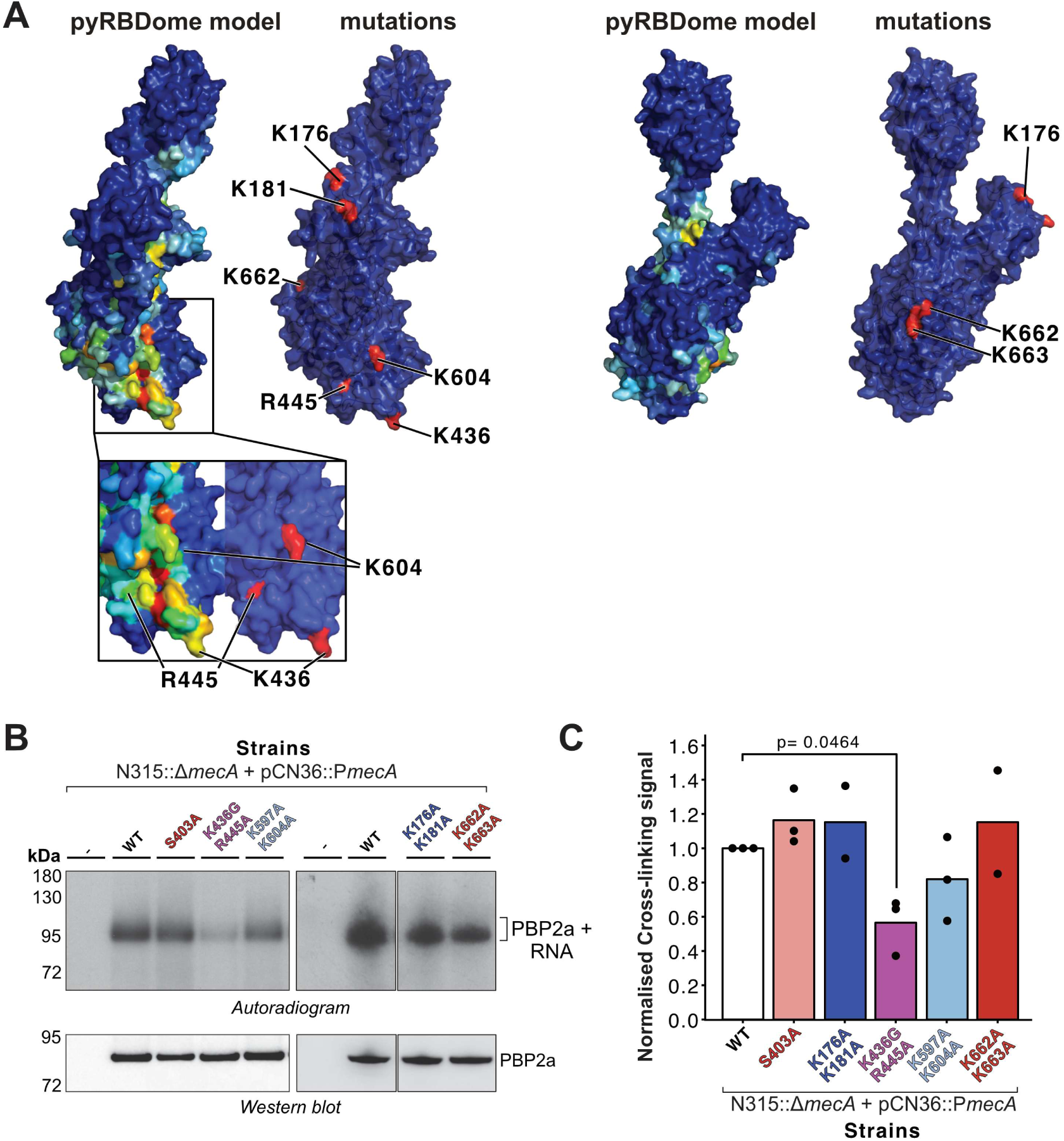
Mutating two positively charged residues in the predicted RNA-binding cleft significantly reduces RNA-binding *in vivo*. **A)** Structure of PBP2a highlighting the pyRBDome prediction results and the amino acids that were mutated. The black box highlights the transpeptidase domain. **B)** The K436G-R445A mutant is RNA-binding deficient *in vivo*. CRAC was performed with N315 *mecA* knockout strains complemented with plasmids expressing the various mutants. WT indicates the wild-type PBP2a protein. Autoradiography was performed to detect cross-linked RNA and Western blot analysis (anti-TAP antibody) was performed to detect PBP2a. **C)** Quantification of the CRAC results in (B). Two to three independent CRAC experiments were performed with each mutant. Autoradiography signals from the CRAC experiments were normalised to corresponding Western blotting signals and compared relative to the results from WT PBP2a (set to 1).

Two to three independent CRAC experiments were performed to assess the RNA-binding capacity of these mutants (Figure 7B-C). Western blot analysis was performed to measure the amount of protein purified during the CRAC purification experiments to normalise for differences in protein recovery (Figure 7B). Quantification of the cross-linking signals (autoradiogram in Figure 7B) normalised to protein levels (Western blot in Figure 7B) revealed that only the K436G-R445A mutations reproducibly caused a substantial reduction in PBP2a binding to RNA *in vivo* (Figure 7C) relative to the parental strain. The strong impact of K436G-R445A on RNA-binding indicates that PBP2a binding to RNA is dominated by electrostatic interactions with the RNA backbone. The K597A-K604A mutation appeared to slightly reduce binding in some experiments, but the results were not statistically significant (Figure 7C). The PBP2a constructs with mutations outside of the predicted RNA-binding region (K176A-K181A and K662A-K663A) showed comparable cross-linking efficiencies as the wild-type PBP2a (Figure 7B-C).

We also performed additional EMSAs to compare the RNA binding of wild-type recombinant PBP2a with two recombinant PBP2a mutants (Supplementary Figure 4A) that showed strong (K436G-R445A) to modest (K597A-K604A) reductions in RNA-binding activity *in vivo*. For this purpose, we used the Hu RNA that cross-linked to PBP2a in the CRAC experiments as well as the corresponding DNA sequence, to determine if the protein discriminates between the two types of nucleic acids (Supplementary Figure 4B-D). This revealed that both mutants were still capable of binding RNA and DNA, with similar effectiveness to the WT. Thus, *in vitro*, sufficient electrostatic attraction remains within these mutants for binding RNA.

Given that the predicted RNA-binding residues we mutated were in proximity to the transpeptidase active site, we asked whether these mutations impacted PBP2a TP activity, using oxacillin resistance as a readout for their catalytic activity. The mutations did not cause notable growth defects in rich TSB medium but reduced oxacillin resistance (Figure 4A). Notably, the RNA-binding deficient mutant (K436G-R445A) retained slightly better antibiotic resistance compared to the catalytically dead mutant (Figure 4A), suggesting that RNA binding and catalytic activity are partially separable functions. Finally, none of the mutations in the predicted RNA-binding cleft noticeably altered protein expression levels (Figure 4B).

### PBP2a RNA binding and TPase activities function independently *in vitro*

Our CRAC analysis of PBP2a mutants implied that RNA-binding and TPase activity are in some way interconnected. However, we cannot exclude the possibility that the mutations we introduced altered the folding of the TPase domain and therefore the protein lost some of its catalytic capacity. Therefore, to substantiate these results, we performed several *in vitro* biochemical assays to gain more insights into the relationship between RNA-binding and TP enzymatic activity. Firstly, we tested whether occupying the PBP2a active sites with the antibiotic ceftaroline (Figure 6A) impacts the ability of PBP2a to bind RNA. We preincubated recombinant PBP2a with increasing concentrations of the antibiotic, followed by addition of RNA to monitor formation of protein-RNA complexes. This revealed that even a 16-fold molar excess of ceftaroline relative to PBP2a was not sufficient to block binding of PBP2a to RNA *in vitro* (Supplementary Figure 6A). When PBP2a was incubated with a 32-fold excess of ceftaroline, PBP2a-RNA interactions were no longer detected (Supplementary Figure 6A). However, it is important to note that achieving this ceftaroline concentration required a final DMSO concentration of 10%, which could not be reduced further, and we observed aggregation of PBP2a under these conditions (Supplementary Figure 6A). Therefore, the loss of PBP2a–RNA complex formation observed at 128µM ceftaroline is more likely attributable to DMSO-induced aggregation than to genuine competition by ceftaroline. Importantly, a bocillin competition assay, in which we tested whether pre-incubation with ceftaroline blocked subsequent bocillin binding to the PBP2a TPase site, confirmed that our ceftaroline batch was active (Supplementary Figure 6B).

We next asked whether RNA could influence TPase activity allosterically. We addressed this question using a previously published LC/MS assay with recombinant PBP2a and PBP2_S398G_, Gly5-Lipid II substrate, and an excess of specific RNA or DNA oligonucleotides (see Materials and Methods and Supplementary Table 3). While we confirmed that our recombinant PBP2a was enzymatically active, little cross-linked product was formed in this assay (Supplementary Figure 7A-C). We also tested various RNA and DNA sequences in this assay, sequences that were frequently detected in our CRAC data. Addition of a 5-fold excess of these substrates did not significantly impact dimer formation, suggesting that nucleic acids do not impact the TPase activity of PBP2a (Supplementary Figure 7A-C). However, given the inherently low TPase activity of PBP2a in this reconstituted system, it is certainly possible that any stimulatory or inhibitory effect of nucleic acids on PBP2a activity would fall below the detection limit of our assay.

Taken together, we conclude that it is very unlikely that RNA competes with PBP2a’s binding to its normal Gly5-Lipid II substrate or affects TP activity.

## Discussion

With the rapid development of high-throughput techniques that allow global capture of RNA-binding proteomes (RBPome), recent studies have discovered a plethora of novel RBPs. This includes many metabolic enzymes capable of binding RNA, suggesting a linkage between RNA regulation and metabolism (30). Recent work in our lab characterised the RBPome of MRSA, hinting that many metabolic enzymes, including enzymes involved in cell wall biosynthesis, are putative novel RBPs (2). It remains, however, unclear how many false-positive hits our RBPome approaches produced and therefore validation of these findings using alternative approaches becomes critical.

Much to our surprise, PBPs cross-linked very efficiently to RNA in our MRSA RBPome studies (2). This seemed very counterintuitive, given the cell surface localisation of these proteins. In this study we further characterised the RNA-binding activities of PBP2a, the enzyme that confers resistance to β-lactam antibiotics.

### PBP2a binds hundreds of transcripts, without apparent specificity and without impacting their steady state levels

Our CRAC experiments showed that PBP2a cross-links to hundreds of transcripts across diverse classes. However, when we compared the CRAC data to RNA-seq data generated from the same samples, only a handful of transcripts were significantly enriched in the cross-linking studies. Moreover, binding of PBP2a was also detected throughout many transcripts, indicating that the protein does not recognise any specific sequence or structural motif. Traditionally, RNA binding is associated with co- and post-transcriptional regulation of the RNA substrate. However, while PBP2a binds RNA robustly *in vitro* and *in vivo*, deletion of the *mecA* gene, which encodes PBP2a, did not reveal any changes in RNA steady-state levels compared to the parental strain. These data provide strong evidence that PBP2a does not act as a classical post-transcriptional regulator.

The lack of apparent specificity in RNA binding and the observation that PBP2a associates with a large portion of the transcriptome made mechanistic interpretation of these findings very challenging. One alternative explanation is that PBP2a resides in a negatively charged environment (due to teichoic acids and peptidoglycans) and therefore might have at least some affinity for negatively charged molecules such as RNA. One could argue that this could lead to non-specific capture of proximal RNA during UV cross-linking. However, several lines of evidence argue against this “proximity capture” model. First, a recent study showed that π-stacking between aromatic residues and uracil residues initiates UV cross-linking of proteins to RNA (31), indicating that successful cross-linking requires specific molecular interactions rather than simple proximity. Secondly, it was striking that mutating only two positively charged residues predicted to bind RNA (Figure 7) was sufficient to substantially reduce UV cross-linking to RNA *in vivo*. Collectively, these findings strongly argue against “chance binding” of RNA to PBP2a and suggest *bona fide* RNA-binding activity.

### What is the function of RNA binding to PBP2a?

The obvious question that remains is therefore, what is the biological significance of PBP2a binding? Based on the work presented here as well as previously published work (32, 33), we favour a model where RNA aids the folding, translocation and oligomerisation of PBP2a. RNA might act as a scaffold or modulator of protein folding and protein-protein interactions during PBP2a folding. The PBP2a mutant lacking the N-terminal transmembrane region *(Δmem*) bound RNA even more robustly (Figure 5C), hinting that RNA-binding may occur in the cytosol with partially folded or even unfolded PBP2a. Our EMSA results showed that PBP2a oligomerises on RNA *in vitro* (Figure 3), supporting a role for RNA as a scaffold. But demonstrating a scaffolding role for RNA *in vivo* is not trivial. Intriguingly, PBP2a also oligomerises within lipid raft-like membrane microdomains *in vivo* termed functional membranes (32, 33), where the chaperone FloA prevents aggregation of partially folded PBP2a. RNA is likely present in these membrane microdomains, as RNA export to extracellular vesicles (EVs) (34) involves interactions with raft-like structures (35, 36). Notably, RNA can function as a chaperone and collaborate with protein chaperones to facilitate protein folding (37, 38). The final folding stages of PBP2a occur at the cell surface with the assistance of extracellular chaperones (22). Given that (i) PBP2a requires quality control mechanisms to achieve proper folding, (ii) PBP2a oligomerises on RNA *in vitro*, (iii) removing the PBP2a membrane anchor (*Δmem*) increases RNA-binding, suggesting interaction with unfolded states, and (iv) RNA is present at membrane microdomains, it is tempting to speculate that RNA may interact with nascent PBP2a during co-translational insertion and early folding stages at the cytoplasmic face of membrane rafts, potentially preventing premature aggregation alongside FloA, before the fully translocated protein undergoes final maturation by extracellular chaperones (PrsA/HtrA1) at the cell surface.

## Materials and Methods

### Bacterial strains and general culture conditions

All strains in this study are listed in Supplementary Table 1. *S. aureus* N315 was used as a parental strain. *E. coli* DH5α was used for recombinant DNA work and general plasmid propagation, while *E. coli* IM08B - mimicking *S. aureus* adenine methylation patterns (39) was used for propagating plasmids destined for N315. For recombinant protein expression and purification, the *E. coli* T7 Express lysY/Iq strain was used.

*E. coli* and *S. aureus* were grown in lysogeny broth (LB) and tryptic soy broth (TSB), respectively, at 37°C with shaking at 200 rpm, except when using the temperature-sensitive plasmid pIMAY, which requires 30°C for replication. Media were supplemented with antibiotics when appropriate, at the following concentrations: chloramphenicol at 15 μg/mL (*E. coli*) or 10 μg/mL (*S. aureus*); ampicillin at 100 μg/mL (50 μg/mL for the pCN36 vector), tetracycline at 10 μg/mL. To assess MRSA antibiotic susceptibility, oxacillin was used at 0.2, 2, 4, 16 or 64 μg/mL.

### *S. aureus* electrocompetent cells preparation and electroporation

A single colony of *S. aureus* N315 was grown overnight and then diluted to OD_595_ 0.05. When OD_595_ reached 0.4-0.6, the culture was chilled on ice for 10 min, and cells were harvested by centrifugation at 3,220 x g for 10 min at 4°C. Cell pellets were washed with one volume of cold 0.5 M sucrose, resuspended in half-volume of cold 0.5 M sucrose and incubated on ice for 30 min. Cells were then pelleted again and resuspended in 1:200 volume cold 0.5 M sucrose. Aliquots (45 μL) were flash-frozen in liquid nitrogen and stored at −80 °C or instantly used for electroporation.

For electroporation, 45 μL of thawed electrocompetent cells were mixed with 1-5 μL DNA and incubated on ice for 5 min. Cells were transferred to a chilled 0.1 cm cuvette and electroporated at 2.1 kV, 200 Ω, 25 μF, then recovered in 950 μL TSB for 1 h at 37 °C (or 2 h at 30 °C for pIMAY). Transformants were plated on TSA containing the appropriate antibiotic and incubated at 37 °C overnight or 30 °C for 2 days for pIMAY.

### Deleting and tagging the *S. aureus mecA* gene

Chromosomal tagging of *mecA* with the His_6_-TEV-3xFLAG (HTF) tag and gene deletion were performed using the temperature-sensitive pIMAY plasmid for allelic exchange, involving chromosomal integration and excision through homologous recombination as previously described (2, 40). A pIMAY vector containing the HTF tag, flanked by 1 kb homology arms upstream and downstream of the *mecA* stop codon was constructed using Gibson assembly. Homology arms were PCR-amplified from the *S. aureus* genome, using Q5® High-Fidelity DNA Polymerase (NEB, M0491). For *mecA* deletion, 1 kb homology arms flanking the start and stop codons were cloned into pIMAY. All DNA fragments and primers used are listed in Supplementary table X and Supplementary table X. Recombinant plasmid sequences were confirmed via Sanger sequencing. All pIMAY plasmids were concentrated to at least 1 μg/μL and 5 μL of plasmid was used for electroporating *S. aureus*. pIMAY presence in *S. aureus* was confirmed by colony PCR with the MCS_F/MCS_R primers. Chromosomal integration of the plasmid was induced by incubation at 37°C with chloramphenicol: A single colony of *S. aureus* carrying the plasmid was resuspended in 200 μL of TSB, diluted 1:1000 and 100 μL plated on TSA containing 10 µg/mL chloramphenicol. After overnight incubation at 37°C, 20-40 colonies were streaked on a fresh TSA-chloramphenicol petri dish and incubated at 37°C overnight again. Integration screening was performed by colony PCR, with two primer sets: PBP2a_chr_F/HTF_PBP2a_R (“upstream PCR”) and HTF_PBP2a_F/PBP2a_chr_R (“downstream PCR”). Successful integration yielded a 1100–1200 bp band with either reaction (Supplementary Table 2). Plasmid excision was induced by incubation at 30°C with 1 mg/L anhydrotetracycline (ATc): A single colony confirmed positive for integration was grown overnight at 30°C without antibiotic, then plated (10^−6^ and 10^−8^ dilutions) on TSA with ATc and incubated at 30°C for 2 days. To screen for colonies that had lost the plasmid, 200 colonies were replica-plated on TSA with and without 10 mg/L chloramphenicol and grown at 30°C for 2 days. Chloramphenicol-sensitive colonies were screened through colony PCR with primers PBP2a_chr_F/PBP2a_chr_R and compared with wild-type genomic DNA to confirm *mecA* tagging/deletion. Wild-type S. aureus genomic DNA was used as a PCR control. If chloramphenicol-sensitive colonies were not observed, the process was repeated with fresh TSA petri dishes supplemented with ATc. Successful HTF-tagging or gene deletion was confirmed by Sanger sequencing of the chromosomal region surrounding the integration site.

### Western blot analyses

Proteins resolved by SDS-PAGE were transferred onto a nitrocellulose membrane (Thermo Scientific, 88018), using the iBlot 3 Western Blot Transfer System (Invitrogen). Then the membrane was blocked in a blocking solution containing 5% skim milk powder and 0.1% Tween-20 in phosphate-buffered saline (PBS), shaking for 60 min at room temperature. Subsequently, the membrane was probed with a primary antibody, diluted 1:5000 in blocking solution, rotating overnight at 4°C. Primary antibodies used for probing included anti-TAP (Invitrogen, CAB1001), anti-FLAG (Sigma-Aldrich, F4049), and anti-enolase (Abcam, ab229919). Membranes were washed three times with PBS containing 0.1% Tween-20 (PBS-Tween) for 10 min each and then probed with secondary horseradish peroxidase (HRP)-linked anti-Rabbit antibody (Invitrogen, A16096), diluted 1:5000 in blocking solution, shaking for 60 min at room temperature. After another 3 washes in PBS-Tween, the protein of interest was detected by chemiluminescence using the Pierce ECL Western Blotting Substrate (Thermo Scientific, 32106), following the manufacturer’s instructions. Proteins were detected using either X-ray films (Thermo Scientific, 34089) or the ImageQuant 800 system (Amersham).

### Complementation of N315::Δ*mecA* strain using pCN36

The *pCN36* staphylococcal shuttle vector was used to reintroduce *mecA* into the N315::Δ*mecA* strain. To construct pCN36::PmecA-*mecA*-HTF, the endogenous *mecA* promoter (P*mecA*) and the *mecA*-HTF fragment were PCR-amplified from the N315::*mecA*-HTF genome using Q5 DNA polymerase with primers N73/N67 (Supplementary Table 2), which included KpnI and EcoRI sites. The PCR product was resolved on a 1% TBE agarose gel, gel-purified, digested with KpnI and EcoRI, and ligated into similarly digested pCN36 (41). Following Sanger sequence verification, the recombinant plasmid was transformed into *E. coli* IM08B for propagation with proper methylation and then introduced into Δ*mecA* electrocompetent cells. Positive clones were detected through colony PCR using the pCN33_F/pCN33_R primers (Supplementary Table 2), and PBP2a-HTF expression was verified through western blotting.

### Site-directed mutagenesis

Mutations were introduced into the *mecA* coding sequence in pCN36::PmecA-*mecA-HTF* using the Q5^®^ Site-Directed Mutagenesis Kit (NEB, E0554S). Mutagenesis primers are listed in (Supplementary Table 2). The kit protocol was followed with the following modifications. For exponential amplification, 40 ng plasmid DNA was used in a 25 μL PCR containing Q5 Hot Start High-Fidelity 1x Master Mix and 0.5 μM of each primer. Thermocycling conditions were: 98°C for 30 s; 30 cycles of 98°C for 10 s, 50-72°C for 30 s, 72°C for 5 min; final extension at 72°C for 2 min. The PCR product was resolved on a 1% TBE agarose gel, and the plasmid band was gel-purified. For the KLD reaction, 80-100 ng of gel-purified DNA was incubated at room temperature for 30 min with 1x KLD buffer and 1x KLD enzyme mix in a 5 μL reaction. The entire reaction was then used to transform DH5α competent cells, which were plated on LB ampicillin 50 μg/mL and grown overnight at 37°C. Positive clones were identified by colony PCR (pCN33_F/pCN33_R), and mutations were confirmed by Sanger sequencing. Verified plasmids were then passed through *E. coli* IM08B and introduced into N315::*ΔmecA* by electroporation.

### *S. aureus* growth curves

To quantify growth rates and determine minimum inhibitory concentrations (MIC) of β-lactam antibiotics in *S. aureus*, the broth dilution method was used (42) in which equal volumes of inoculum and antibiotic are mixed. An overnight culture was diluted in TSB to OD_595_ of 0.1, while antibiotic stocks were diluted in TSB to twice the final required concentration. Then, 100 μL of inoculum and 100 μL of antibiotic were mixed and dispensed into a transparent Greiner 96-well flat-bottom plate. Plates were incubated in an Infinite 200 Pro plate reader (Tecan), shaking at 36.5–37.5°C (535 s duration, 6 mm amplitude) for 16-20 h, with OD_595_ recorded every 9 min. Each experiment included four technical replicates.

Growth curves were generated using the Omniplate Python module (43). OD values were corrected using the correct media function, which subtracts medium-only controls from sample wells. Corrected OD values from the four replicates were plotted over time, with standard deviations shown as shaded errors.

### RNA extraction

RNA extraction from *S. aureus* was performed as previously described (2), with minor modifications. Briefly, 5 mL of culture at OD_595_ ∼ 3.0 was harvested by centrifugation at 12.186 x g for 5 min at 4°C. Cells were resuspended in 650 µL of guanidium thiocyanate (GTC)-Phenol mix (pH 5.4; 1:1 ratio) and lysed by vortexing for 5 min with 400 µL of zirconia/silica beads (diameter 0.1 mm; BioSpec products, 11079101z), alternating 1 min vortexing with 1 min on ice. The mixture was then incubated at 65°C for 10 min and cooled on ice for 10 min. Phase separation was performed by adding 300 μL of chloroform:isoamyl alcohol (IAA) (24:1) and 80 μL of 100 mM sodium acetate (pH 5.2), followed by vigorous vortexing. After 5 min of centrifugation at 12,045 × g, the upper phase containing RNA was transferred in a new 1.5 mL microcentrifuge tube and 500 μL of acid phenol:chloroform:IAA (25:24:1 ratio) was added, followed by vigorous vortexing for 30 s - 1 min. After 2 min of centrifugation at 12.045 × g, the previous step was repeated, this time adding 500 μL of chloroform:IAA to the upper phase. RNA was then precipitated with 2.5 volumes of ice-cold 96% ethanol and 20 μg of glycogen at −80°C for 1 h. RNA was pelleted by centrifugation at 12,045 x g for 30 min at 4°C, washed with 1 mL of ice-cold 70% ethanol, air-dried and resuspended in DEPC-treated water. RNA concentration was measured using the Qubit4 system using the Qubit RNA HS or Qubit RNA BR Assay kits (Invitrogen, Q32852 or Q10210).

### Prediction of PBP2a RNA-binding residues

For the computational analyses of PBP2a structure, the structure of PBP2a in complex with ceftaroline (rcsb ID 3ZFZ) was used. The pyRBDome analyses on PBP2a were performed as previously described (23). To substantiate these analyses, we also docked UCUU RNA sequences (as U’s and C’s preferentially UV cross-link) using the HDOCK webserver with default parameters (28). Docking of the Teg104 RNA fragment onto PBP2a was performed using AlphaFold3 (29) on the AlphaFold server (https://alphafoldserver.com/welcome) using default settings.

### CRAC

The cross-linking and analysis of cDNAs (CRAC) experiments were performed essentially as previously described (2). Slight modifications were introduced in the above protocol to improve cross-linking yields. Firstly, for the strains expressing the endogenously HTF-tagged PBP2a, oxacillin was added to the medium at 0.2 mg/L to boost expression of the protein. Moreover, we found that UV cross-linking can sometimes be less efficient in turbid cell cultures grown in a dark medium, such as TSB. Hence, prior to cross-linking, cells were rapidly transferred to the colourless low phosphate, low magnesium-containing medium (LPM) (2). For the N315::mecA-HTF CRAC experiments, OD595 ∼ 3.0 cultures were shifted from TSB to LPM (5 mM KCl, 7.5 mM (NH4)2SO4, 0.5 mM K2SO4, 8 μM MgCl2, 1 M KH2PO4, 16 mM Tris-HCl pH 7.8, 0.1% casamino acids, 0.3% glycerol) before UV irradiation. For each sample, 100 mL of culture were filtered, cells were resuspended in 100 mL of pre-warmed LPM and immediately UV-irradiated, as described in the previous paragraph. The same approach was also used for test CRAC experiments of generated strains with PBP2a mutations, but with the TSB medium containing 10 μg/mL tetracycline, to retain the pCN36 plasmids. UV cross-linking of cell cultures was performed using the Vari-X-linker (44, 45) using a UV dose of 1 J/cm^2^ at cell densities of OD_595_ of ∼3.0. To cross-link the *mecA*-HTF and Δ*mecA* strains carrying the pCN36 plasmids, 100 mL of culture was UV irradiated and cells were then harvested on 0.45 μm filters by vacuum filtration (44), filters were flash-frozen in liquid nitrogen in 50 mL tubes and stored at −80°C until use. The resulting data were analysed using the pyCRAC package and the CRAC_PE pipeline as previously described (17, 45)

### RT-qPCR

For RT-qPCR analyses, RNA was extracted from wild-type N315 and Δ*mecA*. To remove DNA contamination, 10 μg of total RNA was treated with 4 U RQ1 DNase in the presence of 4 U Superase-In RNase inhibitor, in a total volume of 15 μL, for 20 min at 37°C. RNA was then purified using RNAClean XP beads (Beckmann Coulter, A63987) and 10 ng was used as input for one step RT-qPCR (Luna Universal One-Step RT-qPCR kit; NEB, E3005). Five µL reactions were run on the LightCycler 480 (Roche). Analysis of the qPCR data was performed using the ΔΔCt method (46).

### Fluorescent labelling and annealing of RNA oligonucleotides

Fluorescently labelled RNA oligonucleotides were used as substrates for Electrophoretic Mobility Shift Assays. The 6FAM-labeled Hu RNA oligonucleotide was purchased from Integrated DNA Technologies (IDT). The Teg104 RNA oligonucleotide was labelled with IRDye 800CW Maleimide (LI-COR Biosciences, 929-80020) using the 5’ EndTag™ DNA/RNA Labeling Kit (Vector Laboratories, MB-9001), according to the manufacturer’s procedures. Both RNA oligonucleotides were folded by incubation at 94°C for 2 min and then cooled to 4°C at the minimal ramp rate of a thermocycler, for 20 min in total.

### LC/MS assay to evaluate PBP2a activity *in vitro*

To evaluate the PBP2a activity *in vitro*, purified PBP2a (2 µM) was incubated at 4°C for 1h with RNA/DNA oligos (10 µM; Supplementary Table 3) in 1x reaction buffer (50 mM HEPES, pH 6.5, 2.5 mM MgCl_2_, 0.05 % DDM). Later, *S. aureus* Gly5-Lipid II in DMSO (40 µM) and PBP2_S398G_ (2 µM) were added and incubated for 3 hours at RT. The reaction was then quenched at 95 °C for 5 min followed by 2 hours of treatment with mutanolysin (from *Streptomyces globisporus*, Sigma, 2 U) at 37 °C. Sodium borohydride (10 µL of 10 mg/mL solution) was used to reduce the resultant disaccharides for 30 min. The pH was then adjusted to 4 with the addition of 2 µL of 20% phosphoric acid. The reaction mixture was lyophilized and then redissolved in 12 µL of H_2_O miliQ and subjected to LC/MS analysis, conducted with Ultra-Performance Liquid Chromatography system interfaced with a Xevo G2/XS Q-TOF mass spectrometer (Waters Corp.). Chromatographic separation was achieved using an ACQUITY UPLC BEH C18 Column (2.1 mm × 150 mm, 1.7 μm pore size. Waters Corp.) heated at 45°C. 0.1% formic acid in Milli-Q water (buffer A) and 0.1% formic acid in acetonitrile (buffer B) were used as eluents. The gradient of buffer B was set as follows: 0–3 min 5%, 3–6 min 5–6.8%, 6–7.5 min 6.8–9%, 7.5–9 min 9–14%, 9–11 min 14–20%, 11–12 min hold at 20% with a flow rate of 0.175 ml/min; 12–12.10 min 20–90%, 12.1–13.5 min hold at 90%, 13.5–13.6 min 90–2%, 13.6–16 min hold at 2% with a flow rate of 0.3 mL/min; and then 16–18 min hold at 2% with a flow rate of 0.25 mL/min. The following ions were extracted from each chromatogram: monomer: 1253.5856 (M+1); and dimer: 1209.0617 ((M+2)/2). The QTOF-MS instrument was operated in positive ionization mode. The collision energy was set to scan between 6 eV and 15–40 eV. Mass spectra were acquired at a speed of 0.25 s/scan, and the scans were in a range of 100–2000 m/z. Data acquisition and processing were performed using UNIFI software package (Waters Corp.).

### Lipid II extraction and LC/MS analysis of delipidated Gly_5_-Lipid II

Lipid II was extracted as previously described in (16) with some modifications. An overnight culture of *S. aureus* RN4220 in TSB media (10-15 mL) was used to inoculate each 1.5 L culture. The cultures were grown at 37 °C with shaking for 3 hours. When OD_595_ reached 0.5-0.6, moenomycin was added at a final concentration of 0.6 μg/mL to the 1.5 L culture to accumulate Lipid II. Before harvesting cell pellets (4000 rpm for 10 minutes at 4 °C), the treated cultures were cultivated for an additional 20 minutes while shaking. The pellets from 1.5 L cultures were resuspended in 15 mL PBS (pH 7.4) and added to a 125 mL Erlenmeyer flask containing a mixture of 52.5 mL CHCl3:MeOH (1:2). The mixture was high-speed stirred for 1 hour at RT to ensure cell lysis. The mixture (about 70 mL) from the Erlenmeyer flask was poured into two Teflon tubes and centrifuged at 4,000x g for 10 min at 4 °C. The supernatant, which contains the solubilized cellular contents, was collected while the cell debris was visible at the tube’s bottom as a pellet. For each two tubes, the supernatants were combined and poured into a clean 125 mL Erlenmeyer flask containing 30 mL CHCl3 and 22.5 mL PBS (pH 7.4). For an hour, the liquid was swirled rapidly to thoroughly combine the layers. The homogenized mixture was quickly poured into three clean Teflon tubes and centrifuged at 4,000x g for 10 min at 4 °C. It is important to quickly transfer the heterogeneous mixture into three tubes so that the composition of the mixture in each tube is roughly the same. An interface fraction was revealed in each Teflon tube between the top aqueous and bottom organic layers. The aqueous layer was first progressively removed with a Pasteur pipette to get the interface fraction. The combined interface was dried on *Speedvac*. In the second extraction (to remove Park nucleotide), the combined dried interface was dissolved in a 15 mL organic mixture of 6M pyridinium acetate: n-butanol (1:2) (6M pyridinium acetate was prepared by mixing 51.5 mL glacial acetic acid with 48.5 mL pyridine), and washed with 15 mL of aqueous solvent (n-butanol saturated water) in a separatory funnel. The aqueous layer was extracted again with 10 mL organic solvent (1:2, 6M pyridinium acetate: n-butanol) to maximize Lipid II extraction. The organic layers were combined and washed with aqueous solvent (n-butanol saturated water) three times (10 mL x3) to remove the water-soluble Park nucleotide. On *Speedvac*, the pure organic layer was concentrated before being redissolved in DMSO.

To remove the Lipid II tail, the sample resuspended in DMSO from the previous step was incubated with 800 µL of water and 100 µL of 0.1 M ammonium acetate pH 4.2. The mixture was boiled at 100°C for 90 min and then dried on *Speedvac*. After drying, it was resuspended in 50 µL of water and centrifuged at 16000 x *g* for 10 min to remove the precipitate. The pH of the supernatant was adjusted to pH 3-4 and the sample was subjected to LC/MS analysis.

### Recombinant PBP2a expression and purification

To produce recombinant PBP2a, the *mecA* coding sequence lacking the transmembrane N-terminal anchor domain (amino acids 25-668) was amplified from *S. aureus* N315 genomic DNA, using Q5 DNA polymerase, with primers N63/N64, containing NcoI and XhoI restriction sites, respectively. Primer N63 also contained a TEV protease cleavage site, enabling removal of N-terminal His6 tag following nickel purification. PCR-amplified DNA was resolved on a 1% TBE agarose gel, the band of interest was gel-purified, digested with NcoI (NEB, R0193) and XhoI (NEB, R0146), and then ligated into a pRSET plasmid, containing the His6 tag, digested with the same restriction enzymes. The recombinant pRSET-His6-TEV-*mecA* plasmid was subsequently transformed to the *E. coli* T7 Express *lysY/Iq* strain and positive clones were selected by colony PCR using the T7_F/T7_R primers. To construct the pRSET-His6-TEV-*mecA*S403A plasmid, the same procedure was followed, but the *mecA* fragment was amplified using the pCN36::P*mecA*-*mecA*S403A-HTF plasmid as template.

To produce recombinant PBP2a (mutant) proteins, E. coli T7 Express lysY/Iq cells carrying the pRSET expression plasmids were grown to OD_595_ 2.0 at 37°C. Protein expression was induced by adding 0.5 mM IPTG, followed by growth at 18°C for 16-20 h. Cells were pelleted by centrifugation (2,500 x g for 30 min at 4°C), in a Sorvall centrifuge, resuspended in 40 mL Buffer A (50 mM Tris-HCl pH 8.0, 300 mM NaCl, 5 mM DTT, 1 mg/ml lysozyme, 1 EDTA-free protease inhibitor cocktail (Roche; mini)), and lysed by sonication at 4°C (10s on/ 50s off, 80% amplitude). The lysate was then centrifuged at 20,000 x g for 30 min at 4°C and the supernatant was collected and passed through a 0.45 μm filter to remove residual cell debris. To purify the recombinant proteins, lysates were rotated with 3 mL of pre-equilibrated Ni-NTA agarose (Qiagen, 30250) for 1h at 4°C. The beads were then loaded in a polypropylene column (Qiagen, 34964) and the flowthrough was discarded. The nickel resin was washed with 300 mL of Buffer B (50 mM Tris-HCl pH 8.0, 500 mM NaCl, 5 mM DTT, 20 mM Imidazole). To elute the His6-tagged proteins, elution buffers were made using Buffer C (50 mM Tris-HCl pH 8.0, 300 mM NaCl, 5 mM DTT) with increasing imidazole concentrations (50 mM, 100 mM, 200 mM, 300 mM and 400 mM). Four mL of these elution buffers were added sequentially to the resin and eluates were collected. The purity of the recombinant proteins in the eluates was assessed by SDS-PAGE and Coomassie brilliant blue staining. The purest fractions were loaded in a dialysis membrane (SnakeSkin™ Dialysis Tubing, Thermo Scientific, 68100) and 1 mg of home-made GST-TEV protease per 50 mg of PBP2a was added in the dialysis membrane to remove the His6 tag. Dialysis was performed in 2 L of dialysis buffer (20 mM HEPES-NaOH, pH 8.0, 150 mM NaCl) overnight at 4°C. The following day, the dialysed solution was incubated with 1 mL Ni-NTA agarose beads (pre-equilibrated with dialysis buffer) for 1h at 4°C to bind the cleaved His6 tag. The flowthrough was subsequently incubated with 150 μL of Glutathione Sepharose resin (GE Healthcare, 71024800-EG) pre-equilibrated with dialysis buffer, to capture the GST-TEV protease. The supernatant was subsequently collected and concentrated using Pierce protein concentrators (Thermo, 88528), by centrifugation at 3,220 x g at 4°C, until the volume of the protein solution reached 0.5 mL - 2 mL. Fifty μL protein aliquots were frozen in liquid nitrogen and stored at - 80°C.

*S. aureus* PBP2 variant containing an inactive transpeptidase domain (PBP2^S398G^) was overexpressed and purified as previously described (47).

### Statistical analyses

Data were analysed using GraphPad Prism, version 8.0, software (GraphPad Inc. San Diego, CA). *t*-test and one-way ANOVA with multiple comparisons were used for data analysis. Significance was established at *P* values ≤ 0.05.

## Data Availability

The PBP2a CRAC and RNA-seq data have been made available on NCBI GEO (accession numbers GSE337491 and GSE337429, respectively)

## Author Contributions

Conceptualisation, methodology, supervision and funding acquisition: S.G., F.C., N.C. Investigation: N.C., H-D.N., P.A-R, M.B., G.T. Visualisation: N.C., H-D.N., M.B., G.T., F.C., and S.G. Writing - original draft: N.C. and S.G. Writing - review and editing: all authors.

## Declaration of generative AI and AI-assisted technologies in the manuscript preparation process

During the preparation of this work the authors used Claude Sonnet to correct spelling and grammar mistakes and improve the clarity of the text. After using this tool, the authors reviewed and edited the content as needed and take full responsibility for the content of the published article.

## Acknowledgements

We would like to thank Svetlana Chabelskaia for providing pCN36 vector, Ross Fitzgerald for providing the MRSA N315 strain, Wei Li (李伟) for helping produce recombinant PBP2a and providing reagents, Liang Cui-Chu for helping with analysis of the RBPome data, and Henrik Strahl, Waldemar Vollmer and Víctor M. Hernández-Rocamora for fruitful discussions and helpful feedback. F.C. was supported by Swedish Research Council grants (VR2023-02263), and the Knut and Alice Wallenberg Foundation (KAW2023.0346). G.T. was supported by a postdoctoral fellowship from the Svenska Sällskapet för Medicinsk Forskning (SSMF). S.G. was supported by a Medical Research Council Non-Clinical Senior Research Fellowship (MR/R008205/1) and Medical Research Council Programme grant (MR/Y013131/1). N.C. was supported by a Darwin Trust of Edinburgh doctoral training fellowship.

